# Diversity of Secretion System Apparatus in Tomato Wilt Causing *Ralstonia solanacearum* Strains: a Comparative Analysis Using *in-silico* Approach

**DOI:** 10.1101/2022.04.21.489029

**Authors:** Goutam Banerjee, Sandipan Chatterjee, Pratik Banerjee, Pritam Chattopadhyay

## Abstract

*Ralstonia solanacearum* (Rs) species is the leading cause of bacterial wilt disease in a wide range of host plants worldwide. In the present study, secretion system analysis of five tomato pathogenic Rs strains was carried out *in-silico*. This paper describes a new protocol to identify the secretion system components i.e. SSCs (T1SS-T6SS, Flg, T4P, and Tad-Tat). A total of 865 SSCs were identified using the new protocol. Contributions of SSCs into core-secretion system apparatus (i.e. SSA) were also studied. Synteny was discovered among the secretion system apparatus (SSA) where relative frequency of SSCs to core-SSA is high (>20%) which includes T1SS, T2SS, T5SS, T4P, and Tad-Tat, but excludes T3SS, T4SS, and Flg. To the best of our knowledge, this is the first report indicating that during the evolution of *Rs*, most of the secretion system apparatus (T1SS, T2SS, T5SS, T4P, and Tad-Tat) were highly conserved and came from a single ancestor, while T3SS and T6SS may have arrived later, probably from horizontal gene transfer.

## Introduction

*Ralstonia solanacearum* (hereafter referred to as ‘Rs’) is an aerobic, non-spore-forming, gram-negative, plant pathogenic bacterial species that includes a group of β-proteo-bacteria (1). Rs is considered a major plant pathogen and has been divided into four distinct monophyletic phylotypes (2). Rs exhibits some unique features of plant pathogens, including abundance in rhizospheric soil, large host specificity (>200 plant species), tissue-specific tropism, invasion of the root system, and multiplication and colonization (> 10^9^ c.f.u. per g fresh weight) in xylem vessels (3). Therefore, Rs is declared as ‘priority plant pathogens’ in many countries in the world. Rs is also classified as ‘quarantine organisms’, ‘bioterrorism’, and ‘double usage agents’ by different regulatory authorities in the USA and Europe (4).

Many bacterial secretion systems have been identified in Rs (5). As per the current understanding, bacterial secretion is made up of a number of specialized systems, such as type I-VI secretion systems (T1SS-T6SS), along with flagella (Flg), type IV pili (T4P), and tight adherence (Tad) secretion systems (6). It is well established that bacterial secretion systems contribute to host specificity. Over decades of intensive research, Rs has become a model system for investigating mechanisms of plant pathogenesis (7). However, the study of secretion systems diversity within Rs is rare.

As a model organism, Rs gives us a unique opportunity to study bacterial secretion system apparatus of five tomato pathogenic strains (Rs GMI1000, Rs CFBP2956, Rs CMR15, Rs FQY_4, and RsPSI07), representing four phylotypes. Here, we have tested the hypothesis that there might be a high degree of variation in the secretion system components (SSC) of Rs due to their ability to infect different types of hosts. With the advent of improved data analysis tools, many bioinformatic platforms have been introduced to identify bacterial secretion systems across entire genomes. Previously, secretion systems of 2643 bacterial genome have been investigated by a standalone programme i.e. TXSScan **(6)**. However, in this present *in-silico* investigation, we utilized the KO (KEGG Orthology) based protocol to identify bacterial secretion system-related candidate genes and protein orthologs. Host specificity may have an impact on bacterial secretin system. To avoid this factor, we have selected and explored the diversity of secretion system only the strains that infect tomatoes. This study is important to determine the infection pattern of a deadly pathogen within a common host, and here we have elucidated the impact of current data analysis tools to enhance scientific knowledge. These secretion-related genes might be a future target to control the infection in the agricultural field.

## Materials and Methods

### Data Mining

This in-silico study was undertaken specifically to compare the bacterial secretion systems of five tomato infecting strains of *R. solanacearum* (Table 1). Fully annotated whole genome sequences of *R. solanacearum* strains infecting tomatoes were searched from publicly available databases. Five such genomes were downloaded from NCBI RefSeq (https://www.ncbi.nlm.nih.gov/genome/genomes/490). The strains were categorized on the basis of original literature (Table S1). Orthologs of secretion system components were derived from KEGG-KO (KO or KEGG ORTHOLOGY database of Kyoto Encyclopaedia of Genes and Genomes) database. These orthologs were verified within the selected genomes using KEGG-Genome database.

**Table 1.**
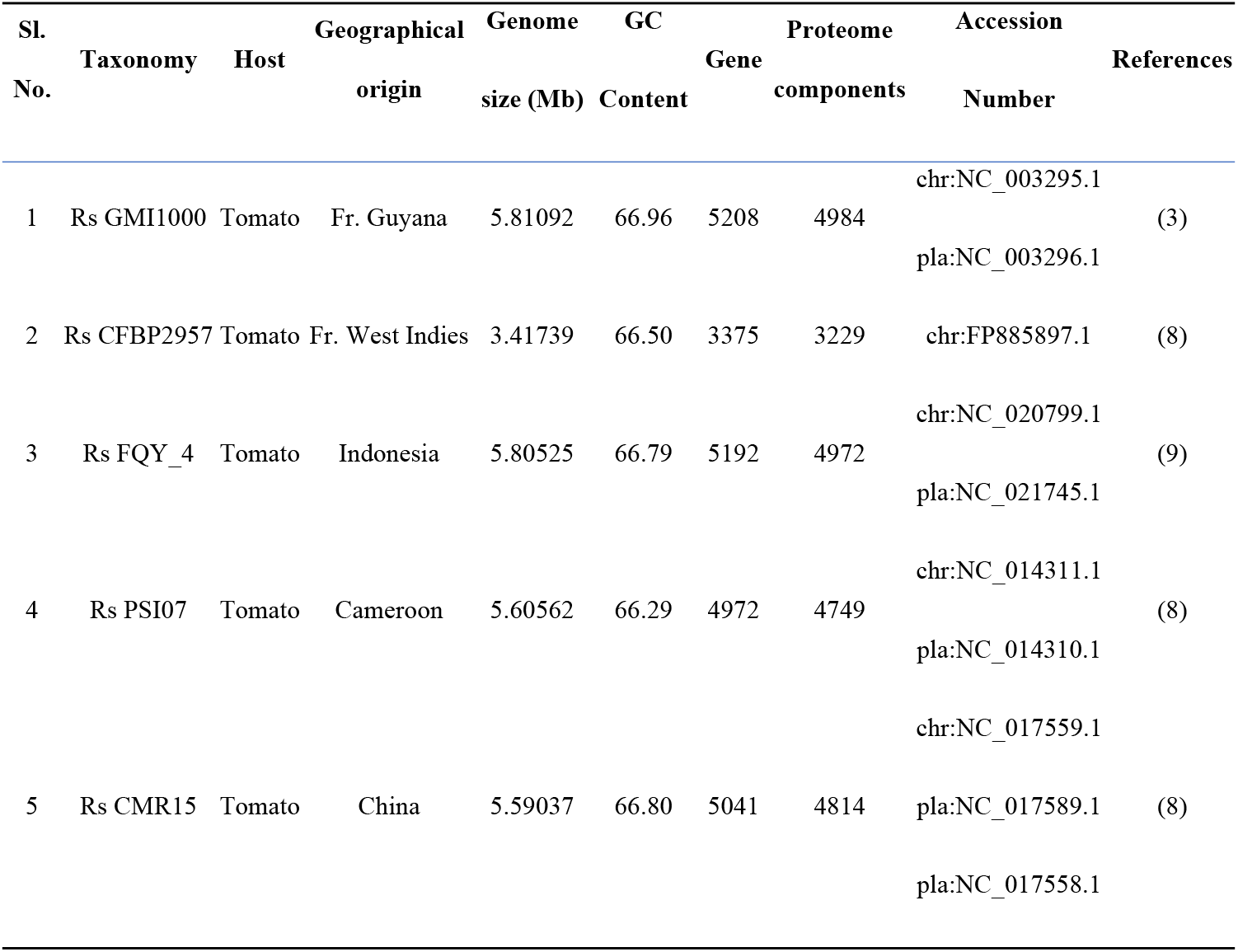
List of the five-tomato pathogenic *Ralstonia solanacearum* genomes used in this study.

### Secretion System Comparison

In order to compare the bacterial secretion systems (T1SS-T6SS, T9SS, Flg, T4P, and Tad), a dataset was prepared from the previously published data by Ma et al. (10), Henderson et al. (11), Desvaux et al. (12), Abby et al. (6), and Lasica et al. (13), categorically presented in Table S1 (considered as reference set). These systems were selected in order to maximize sequence diversity. To identify bacterial secretion system related genes this reference dataset was examined manually using the KO (KEGG ORTHOLOGY) database (Table S2). All the KO (orthologs) was validated by their presence or absence in all the selected R. solanacearum genomes in KEGG-Genome database (Table S3). A flowchart of the experimental design is presented in Fig.1.

**Fig 1.**
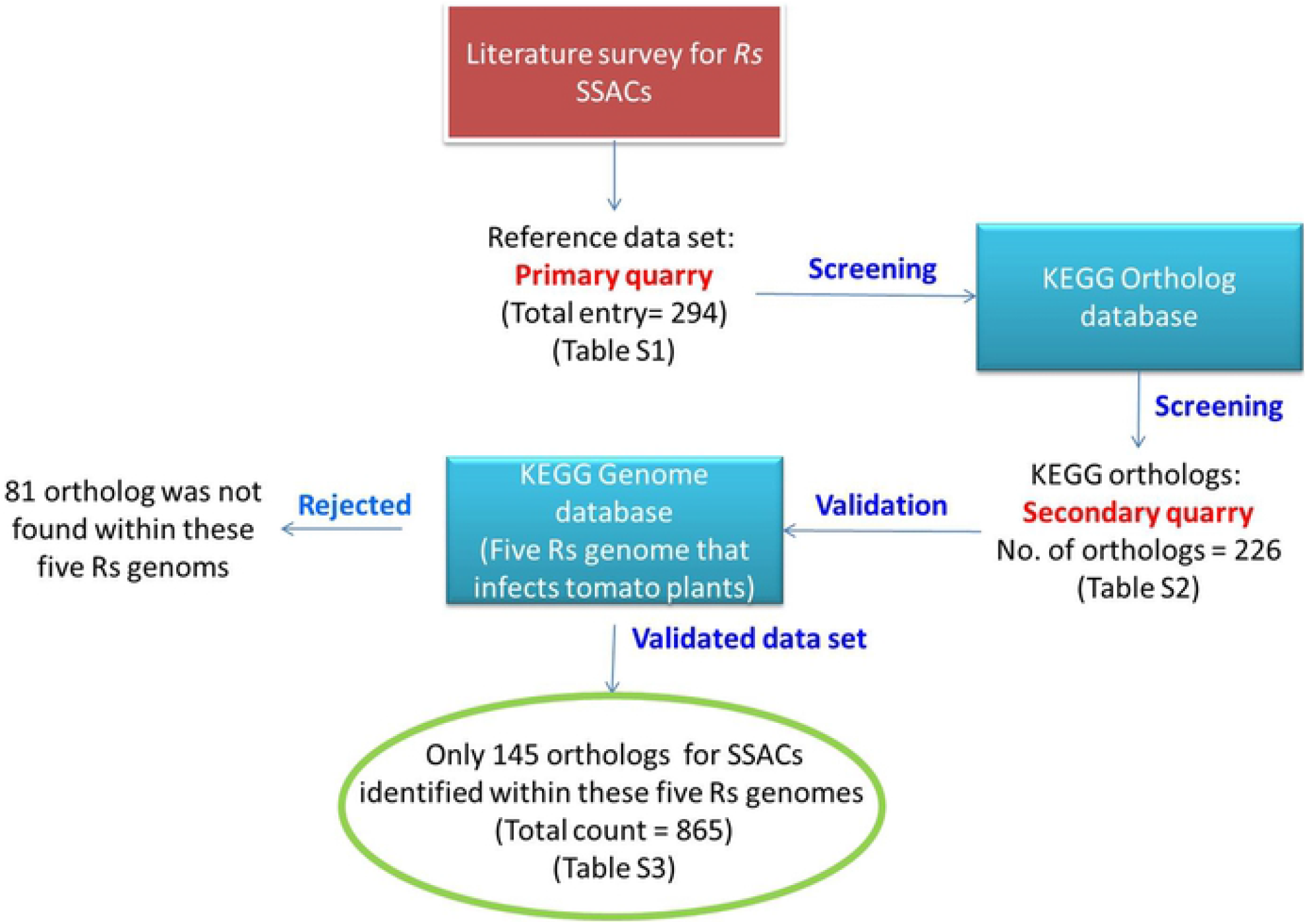
Protocol to identify secretion system components of five tomato pathogenic *R. solanacearum* strains.

### Synteny Analysis

Synteny is the study of the conservation of gene order. It is an important tool to predict the functional relationships between genes and to assess the orthology of genomic regions (14). In this study, SyntTax (a web service designed to take full advantage of the large amount of archaeal and bacterial genomes by linking them through taxonomic relationships) was used to determine the synteny among the SSC subcomponents (14). In SyntTax, the synteny methodology is based on the Absynte algorithm (15).

### Phylogenetic Analysis

The phylogenetic analysis of five R. solanacearum strains was performed based on 16S rRNA gene sequence. Full length 16S rRNA gene sequences were retrieved from the whole genome sequence using RNAmmer 1.2 server (16). The phylogenetic tree was constructed by neighbour-joining algorithm using MEGA 6.0 software (17), taking *Xanthomonas oryzae* pv. oryzae KACC 10331 as an out-group member.

### Statistical Analyses

All SSC-related genes were scored based on their copy numbers obtained from the KO database. Based on this matrix, bacterial secretion system components present in all the R. solanacearum genomes were identified as “core secretion system apparatus” or SSA. Contributions of all the systems and sub-systems in the “super SSC” of R. solanacearum were determined and presented in terms of a Venn diagram and occurrence frequency (%). Venn diagrams were calculated with the help of an online tool provided by Bioinformatics & Evolutionary Genomics (available at: http://bioinformatics.psb.ugent.be/webtools/Venn/). Contribution of the individual secretion system into the “core SSC” was measured through relative frequency. The genetic relationships among the *R. solanacearum* strains were estimated using the Euclidean distance matrix using Past3.17 software (18). The matrix data were further evaluated by Principal Component Analysis (PCA) as minimum distance, using Past 4.08 software. Genetic distance was estimated based on pair-wise comparisons and was represented in the form of a dendrogram (Figure 6). Boot strap analysis was carried out for 10,000 replications of the dendrogram.

## Results

### Identification & distribution of Secretion System Components (SSCs)

Identification of SSC were accomplished in three steps: identification of references from published literature and databases (Total entry 294, Table S1), then searching for the orthologs in KEGG-KO database (Total entry 226, Table S2), and then validation of datasets by KO numbers in KEGG Genome database (Total ortholog entry validated 145, Table S3). In this method, a super *R. solanacearum* SSC (cumulative no. 865) containing nine secretion systems and 145 orthologs (Table S3) were identified (Fig. 1). Furthermore, the unique distribution of SSC into five strains was demonstrated in a matrix plot with protein orthologs (Fig. 2). Among the 145 orthologs, 60 were identified in all the strains and defined as the “core SSC” (Fig. 3a).

**Fig 2.**
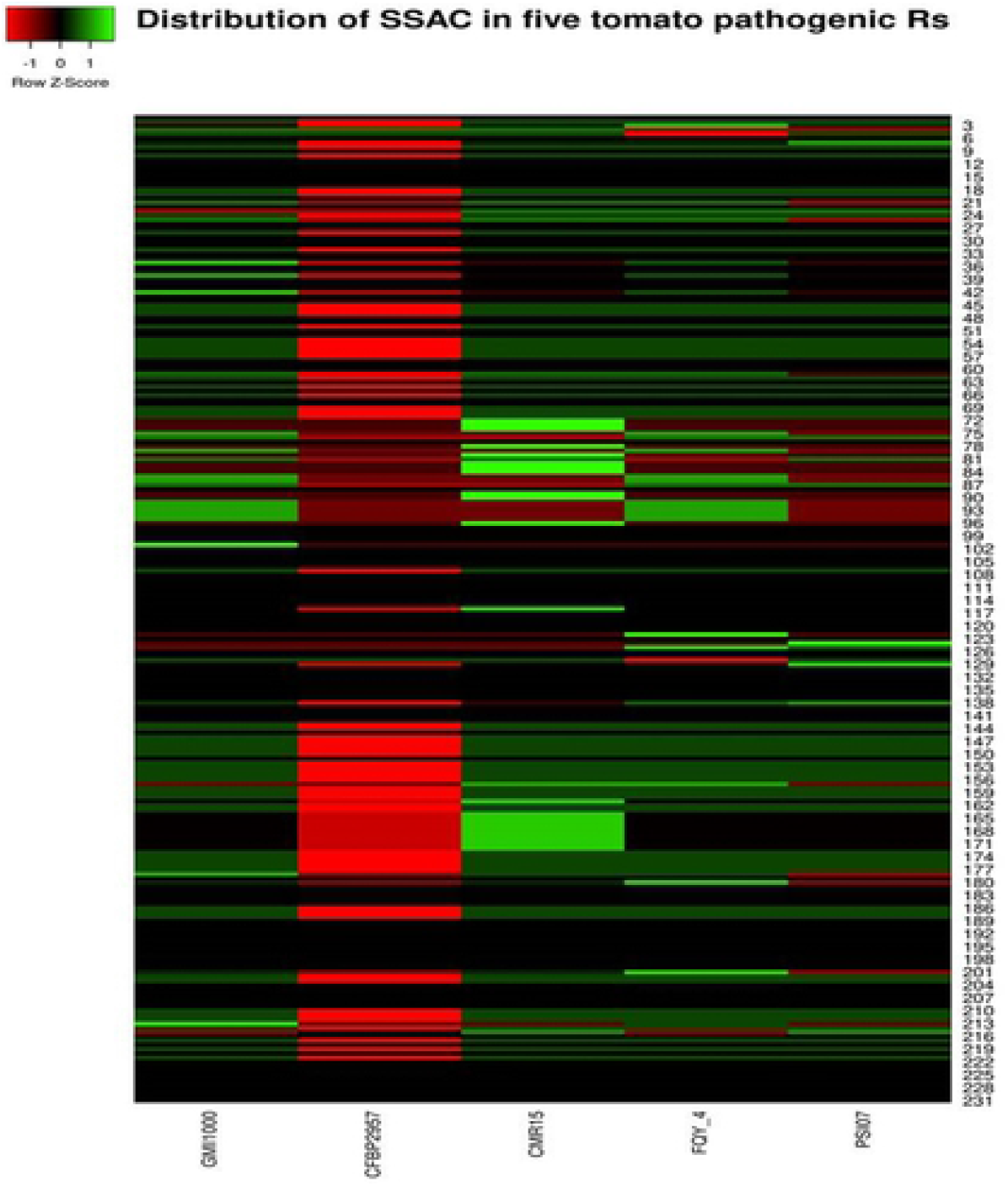
Matrix plot showing similarity and dissimilarity of bacterial secretion systems of five tomato pathogenic *R. solanacearum* strains: The X-axis representing protein orthologs and the Y-axis bars representing strains. Interestingly none of the bars are exact match to anyone. This indicates the individuality of bacterial secretion systems of each strain.

**Fig 3.**
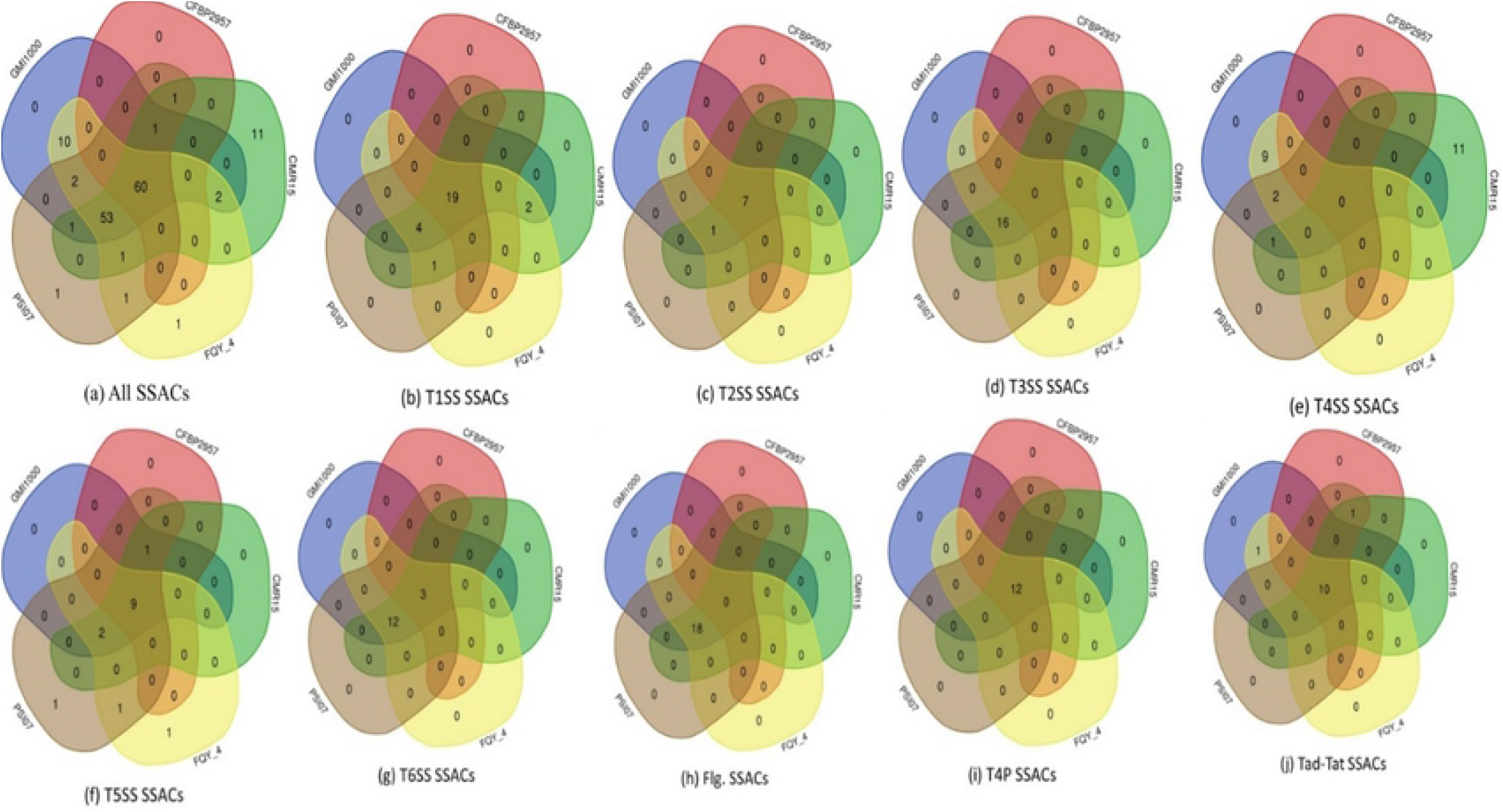
Venn diagrams showing contribution of SSCs into the core-SSC and pan-SSC of five tomato pathogenic *R. solanacearum* strains. **a)** Venn diagrams showing contribution of all SSCs together: Out of 226 orthologs, only 60 were contributed into the core-SSC. **(b)** Venn diagrams showing contribution of T1SS SSACs: Out of 26 orthologs, 19 were contributed into the core-SSC. **(c)** Venn diagrams showing contribution of T2SS SSACs: Out of 8 orthologs, 7 were contributed into the core-SSC. **(d)** Venn diagrams showing contribution of T3SS SSCs: Out of 16 orthologs, none was contributed into the core-SSA. **(e)** Venn diagrams showing contribution of T4SS SSCs: Out of 23 orthologs, none was contributed into the core-SSA. **(f)** Venn diagrams showing contribution of T5SS SSACs: Out of 15 orthologs, 9 were contributed into the core-SSC. **(g)** Venn diagrams showing contribution of T6SS SSACs: Out of 15 orthologs, only 3 were contributed into the core-SSC. **(h)** Venn diagrams showing contribution of Flg SSCs: Out of 18 orthologs, none were not found in Rs CFBP2957. **(i)** Venn diagrams showing contribution of T4P SSACs: Out of 12 orthologs, all were contributed into the core-SSA. **(j)** Venn diagrams showing contribution of Tat-Tad SSCs: Out of 12 orthologs, 10 were contributed into the core-SSC. These diagrams were generated from a web-based application available at http://bioinformatics.psb.ugent.be/webtools/Venn/.

### Type Secretion Systems (T1SS-T6SS)

Nine secretion systems (T1SS, T2SS, T3SS, T4SS, T5SS, T6SS, Flg, T4P and Tat-Tad) were determined in the selected five Rs genomes (Table 2).In this analysis, six type secretion systems were identified (T1SS-T6SS) in all the *R. solanacearum* genomes. However, T9SS was not detected in any of the selected strains as it is specific to the Bacteroidetes phylum (6). In the present investigation, two different T4SS apparatus were identified. One of T4SS secretion system is plasmid-encoded, containing genes for different Vir proteins, and is present only in Rs CMR15. The other T4SS is chromosome encoded and contains different Trb proteins in Rs GMI1000 and Rs FQY_4 (Table S3). We also identified 26 orthologs for T1SS (Fig 3b), 8 for T2SS (Fig. 3c), 16 for T3SS (Fig. 3d), 23 for T4SS (Fig. 3e), 15 for T5SS (Fig. 3f) and 15 for T6SS (Fig. 3g). There is no contribution form the part of T3SS, and T4SS to the core-SSA, and contribution form the part of T6SS is also very low (20.0%). Among type secretion systems, the contribution of T2SS in ‘core-SSA’ was maximum (relative frequency of 87.5%), followed by T1SS (73.08%), T6SS (23.07%), T4SS (22.22%), and T3SS (7.69%) (Fig. 4a). Interestingly synteny of SSCs in T1SS, T2SS, and T5SS, were also observed (Fig. 4b-d). However, synteny was not observed among SSCs of T3SS, T4SS, and T6SS.

**Table 2.** Comparative analysis of bacterial secretion system components (SSC) in five tomato pathogenic strains of *Ralstonia solanacearum*.

| Secretion<br>systems | No. of SSC |  |  |  |  |
| --- | --- | --- | --- | --- | --- |
|  | Rs GMI1000 | Rs CFBP2957 | Rs CMR15 | Rs FQY_4 | Rs PSI07 |
| T1SS | 53 | 33 | 55 | 54 | 53 |
| T2SS | 17 | 07 | 11 | 14 | 11 |
| T3SS | 17 | 00 | 17 | 17 | 16 |
| T4SS | 12 | 00 | 12 | 11 | 03 |
| T5SS | 15 | 11 | 15 | 17 | 19 |
| T6SS | 21 | 03 | 20 | 23 | 23 |
| Flg | 18 | 00 | 28 | 18 | 18 |
| T4P | 26 | 18 | 25 | 26 | 22 |
| Tad | 16 | 08 | 16 | 15 | 16 |
| Tat | 03 | 03 | 03 | 03 | 03 |
| **Total** | **198** | **83** | **202** | **198** | **184** |

**Fig. 4.**
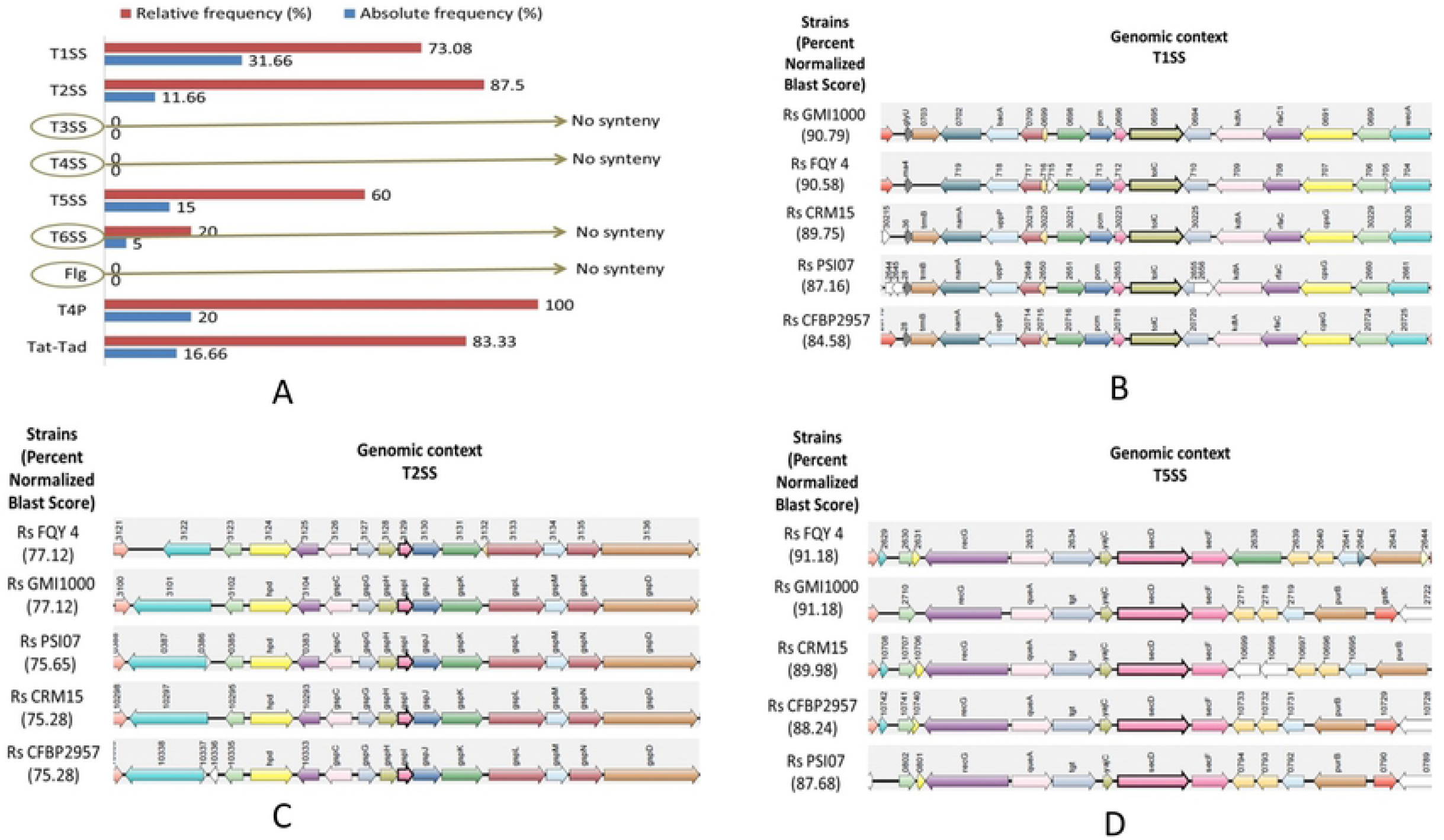
(a) Bar diagram representing contribution of the SSCs into the “core-SSC” of five tomato pathogenic *R. solanacearum* strains: Relative frequency was measured in comparison with particular secretion system, while absolute frequency was measured in comparison with “core-SSC”. Both the frequency was zero for T3SS, T4SS, Flg and very low for T6SS. (b) Synteny diagram showing conservation of T1SS for the five strains infecting same host. (c) Synteny diagram showing conservation of T2SS for the five strains infecting same host. (d) Synteny diagram showing conservation of T5SS for the five strains infecting same host. These diagrams were generated from Synteny webserver.

### Flg

In the present study 18 orthologs (Table S3) for Flg SSACs were identified commonly present in all the experimental genomes except Rs CFBP2957 (Fig. 3h). None of these orthologs have contributed in core-SSA (Fig. 4a). However, no synteny was found for Flg components.

### T4P and Tad-Tat

In the case of T4P SSA, 12 orthologs were identified (Table S3). Among all the SSA, the contribution of T4P in ‘core-SSA’ was maximum (relative frequency of 100%); whereas it is 83.33% for other pili system i.e. Tad-Tat (Fig. 3a). Synteny of T4P and Tad components (Fig. 5a,b) indicate that T4P and Tad originated from a single ancestor.

**Fig. 5.**
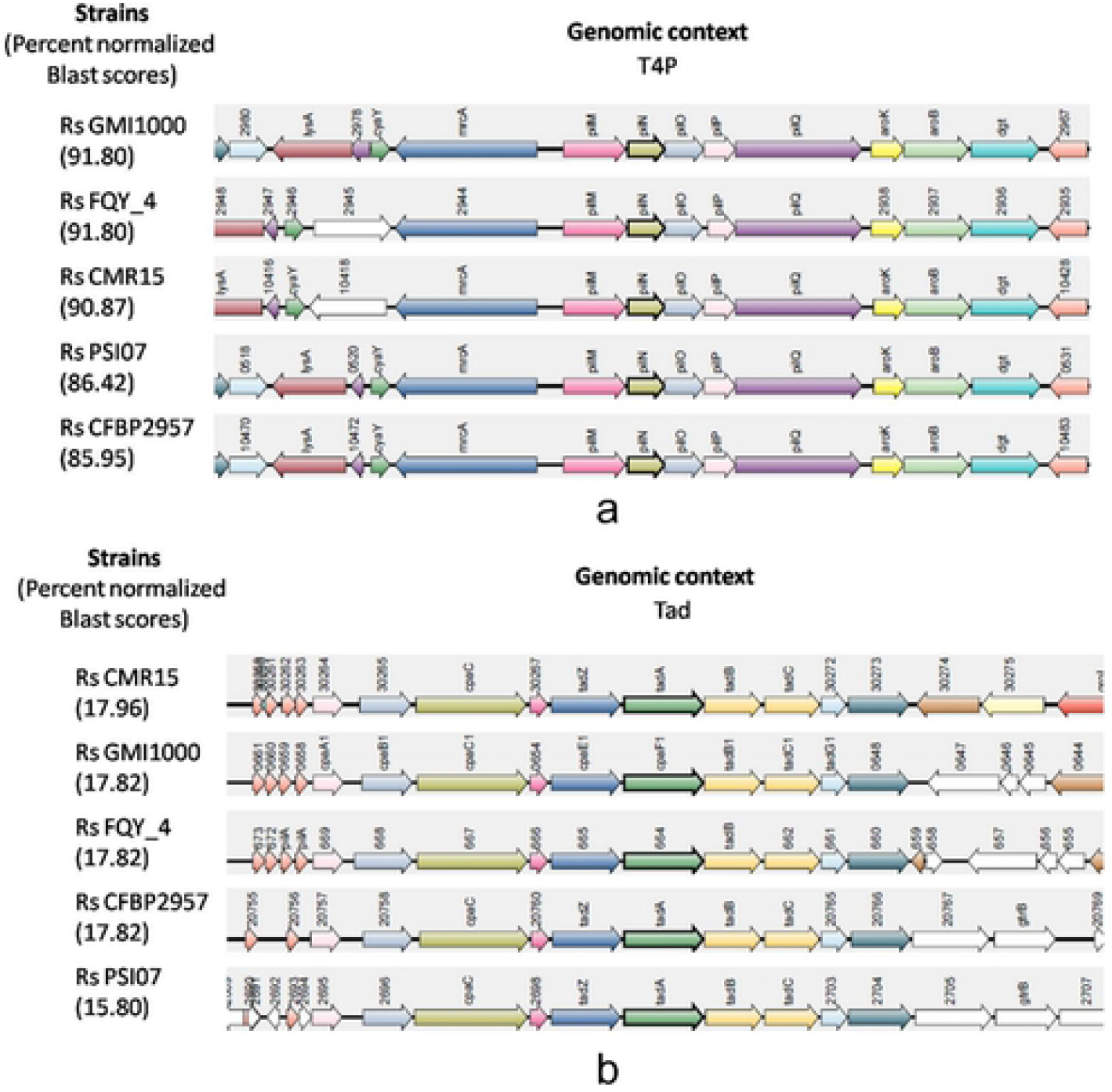
(a) Synteny diagram showing conservation of T4P for the five strains infecting same host. (b) Synteny diagram showing conservation of Tad-Tat for the five strains infecting same host. These diagrams were generated from Synteny webserver.

### Secretion System vs. Phylogeny

Multivariate analysis was conducted based on the similarity matrix drawn from the presence and absence of 145 SSC orthologs. In PCA analysis, three main clusters were identified in the bi-plot comprising PCA1-PCA2 (Fig. 6a), which exhibited differently, clusters of Rs strains. However, the resulting cluster dendrogram (Fig. 6b) showed significant similarity with 16S rRNA based phylogeny (Fig. 6c). Both the clusters dendrogram and phylogenetic tree were supported by high boot strap values (>50) and a common root from the out-group *X. oryzae* pv. oryzae KACC 10331.

**Fig. 6.**
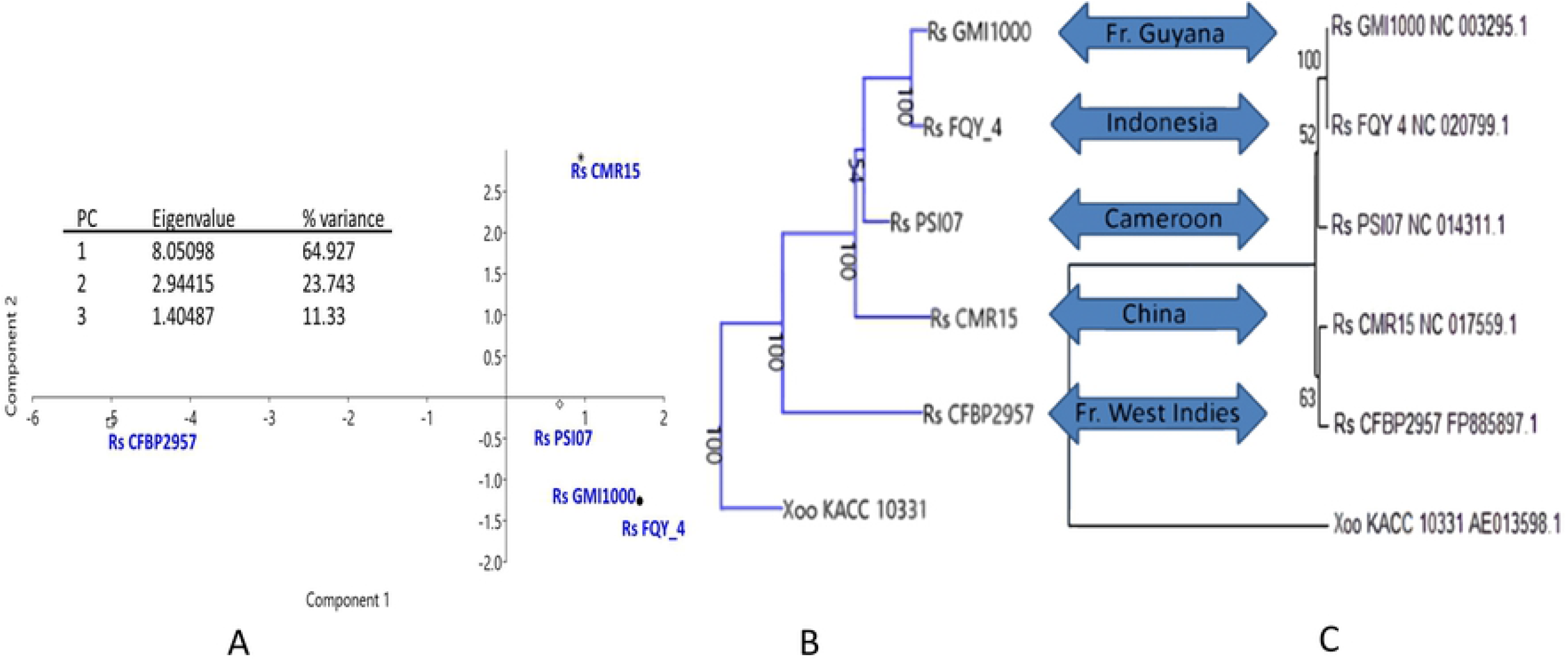
(a) PCA biplot comprising PCA1-PCA2 and shows different clustering of *R. solanacearum* strains on the basis of components of secretion system apparatus. Analysis was done using Past 4.08 software. (b) Cluster dendrogram showing genetic relationships among the five tomato pathogenic *R. solanacearum* strains: Genetic distance was calculated based on pair-wise comparisons using the Euclidean distance matrix based on first two principal components derived from secretion system components (Table S2). (c) Reference phylogenetic tree based on 16S rRNA sequence showing genetic relationships among the five strains of *R. solanacearum:* Genetic distance was calculated based on p-distance, and the phylogenetic tree was constructed by neighbor-joining algorithm using MEGA6 software. Both the cluster dendrogram and phylogenetic tree were compared with the geographical origin of the strains.

## Discussion

In our present investigation, a total of 865 SSC representing 145 orthologs distributed into different recreation system viz. T1SS-T6SS, Flg, T4P and Tad-Tat were identified through the protocol described here (Fig 1). The protocol is based on identification and validation of orthologs from the KO database and Genome database respectively. Annotation of secretion system apparatus components was done earlier by Abby et al. (6). In the present study, manual curating is important to deal with proper annotation of secretion system apparatus components for the following reasons. Firstly, there may be homologous secretion system apparatus components present within a single strain. For example, Vir and Trb proteins of T4SS were homologous. Vir proteins were identified in Rs CMR15, while Trb proteins were identified in Rs GMI100 and Rs FQY_4. Secondly, there may be different orthologs of a single SSC. For example, four orthologs of HlyD were identified (namely: K01993, K02005, K11003, and K12542). Thirdly, even a single ortholog may have more than one copy number within a single strain. For example, GspD (K02453) has four copies in Rs GMI1000. Finally, these copies may be distributed in bacterial chromosomes and plasmids. This may have appeared as a result of gene duplication and then transferred to the plasmid or may have appeared through horizontal gene transfer. Therefore, annotating the SSC orthologs is very critical. Nevertheless, investigation on the SSCs using the KO database has its own limitations. The main limitation is the limited number of genomes available in the KEGG database. In addition to that, there are instances where a KO number was not assigned.

In our present study, all the selected strains were from a single bacterial species (*R. solanacearum)* infecting a common host (tomato). Still, a significant difference was found in their secretion system (Fig. 2). Out of 145 SSC orthologs, only 60 were identified in all the strains and defined as the “core-SSC” for tomato pathogenic strains of Rs (Fig. 3a). The secretion system components have been studied extensively and have repprted in different animal pathogenic bacteria such as *Pseudomonas aeruginosa* (19, 20), *Vibrio cholerae* (21, 22), *Salmonella* sp. (23), *Escherichia coli* (24), *Bacillus subtilis* (25) etc. However, in depth reports regarding the components of secretion system on plant pathogens are scanty. Previously few reports have been published addressing some specific secretion system in *R. solanacearum* such as type III (26) and type VI (27). However, detail observation of all secretion systems in association with host is not reported yet. In this present study, we have explored the distribution of secretion system components and have compared among five RS strains infecting a common host. Result clearly demonstrates that SSCs are intrinsic property of the individual strain, and thus the secretion system of a particular strain is independent of its host.

There are many reports available on the comparative analysis of bacterial strains. But, there are only a few reports on studying the synteny of bacterial secretion systems. The variation and synteny in the T6SS operon within plant and animal associated proteobacteria was previously described by Wu et al. (28). In the present investigation, synteny was observed among T1SS, T2SS, T5SS (Fig. 4b-d), T4P, and Tat-Tad (Fig. 5a,b). Synteny was not observed in T3SS, T6SS, and Flg. Interestingly, T3SS and T6SS are among the lowest contributors to secretory systems for “core bacterial secretion system proteins” (Fig. 4a). This indicates that during the evolution of Rs, most of the secretion systems (T1SS, T2SS, T5SS, T4P, and Tad) were highly conserved and came from a single ancestor. T3SS is responsible for bacterial invasion into the host following intracellular replication of bacteria and triggering apoptosis of the host cell. T6SS is considered an important virulence factor of bacteria which also aids in the formation of biofilms. The absence of synteny in these two systems among the strains of the same bacterial species infecting the same host (tomato) suggests the horizontal transfer of genes in T3SS and T6SS of Rs. (29). To the best of our knowledge, this is the first report on synteny of bacterial secretion systems of tomato wilt causing strains of *R. solanacearum*.

## Conclusions

Our present report provides cluster dendrogram with SSC shows close resemblance with 16S rRNA based phylogeny, which suggests that the bacterial secretion systems are an intrinsic property of the strain. To minimize the secretion system diversity, we selected the Rs strains with a common host; still, we found only 60 out of 145 SSC orthologs as core-SSA. This is for the first time synteny was found to be absent in SSA that contributes less (>20%) in the Core-SSC among the strains of Rs infecting a common host (tomato). To our knowledge, we think this is the first study that clearly indicates that during the evolution of Rs, most of the bacterial SSC (T1SS, T2SS, T5SS, T4P, and Tad-Tat) were highly conserved and came from a single ancestor. T3SS and T6SS have evolved into the strain probably from horizontal gene transfer. Our present findings give a new insight into the Global threat of Rs. T3SS and T6SS components may be targeted to control *R. solanacearum* infestations in the agricultural field in the future.

## Acknowledgments

P.C is very much thankful to Dr. D S Kothari Post-Doctoral Fellowship from University Grant Commission (UGC), Government of India (Award No. F.42/2006 (BSR)/BL/1415/0391 dated 01July, 2015) for providing support. Authors also thanks to the Department of Food Science and Human Nutrition, University of Illinois Urbana-Champaign (USA) for providing bioinformatics facilities.

## Supporting information

**Table S1.** Reference dataset: experimentally validated systems used to define bacterial secretion systems (and related appendages) as a quarry in the KO database.

**Table S2.** Validated dataset: list of the components of bacterial secretion systems (and related appendages) and their orthologs within five tomato pathogenic *Ralstonia solanacearum* strains. Orthologous protein and their locus were identified from KO database (KO no of each protein and multiple locus of each protein was given when available).

**Table S3.** Distribution of SSCs within five tomato pathogenic strains of *Ralstonia solanacearum*

## Author Contributions

**Conceptualization:** Pritam Chattopadhyay

**Data curation:** Sandipan Chatterjee, Goutam Banerjee

**Formal analysis:** Pritam Chattopadhyay, Goutam Banerjee

**Methodology:** Pritam Chattopadhyay, Pratik Banerjee, Goutam Banerjee

**Resources:** Sandipan Chatterjee

**Supervision:** Pratik Banerjee, Pritam Chattopadhyay

**Writing – original draft:** Pritam Chattopadhyay, Goutam Banerjee

**Writing – review & editing:** Pratik Banerjee, Goutam Banerjee, Pritam Chattopadhyay

## Conflicts of Interest

The authors declare no conflict of interest.

## Notes

### Competing Interest Statement

The authors have declared no competing interest.

